# Unveiling the Intricate Sinking behavior of Large Diatoms: Insights from Time-Frequency Analysis of *Palmerina hardmaniana* Sinking under Silicate-Depleted Conditions

**DOI:** 10.1101/2024.03.03.582478

**Authors:** Ping Zhang, Jiachang Lu, Lang Li, Junlian Zhuang, Junxiang Lai, Tianying Shi, Jie Li

**Affiliations:** Guangxi Key Laboratory of Marine Environmental Science, Guangxi Academy of Marine Sciences, Guangxi Academy of Sciences, Nanning, China; School of Resources, Environment and Materials, Guangxi University, Nanning, China

**Keywords:** Sinking behavior, Large diatom, Time-frequency domain, *Palmerina hardmaniana*, Silicate deficiency

## Abstract

Nutrient limits impact diatom sinking in time domain, but response in time-frequency domain is unclear. Studying the response of large diatoms to nutrients exclusively in the time domain fails to fully capture the complete impact of nutrient limitations on sinking behavior due to the absence of crucial information, including period and frequency. Wavelet analysis provides valuable insights into the period and frequency of signals and unveils their positions in the time. This study investigated the sinking behavior response of the large diatom *Palmerina hardmaniana* to silicate-depleted conditions in the time-frequency domain using wavelet analysis. The results showed that *P. hardmaniana* was capable of regulating its sinking speed, and this regulation occurred intermittently in time. The predominant frequency of intermittent regulation fell within the range of 0.13-0.50 Hz (equivalent to a period of 2-8 s) for both control and silicate-depleted conditions. The similarity in the frequency range of regulation between the two groups suggests the involvement of shared physiological mechanisms. *P. hardmaniana* responded to silicate depletion by intensifying the regulation of 0.13-0.50 Hz, which was reflected in the time domain as a change in the proportion of different instantaneous sinking speeds and consequently lead to a significant increase or decrease in the mean sinking speed *(p*<0.05). Additionally, the regulation of *P. hardmaniana* sinking behavior was also influenced by the physiological state of the cells. Short-term silicate stress (30 min) enhanced the oscillation power of sinking regulation, while prolonged silicate stress (≥ 3 d) led to a decline in oscillation power.

## 1. Introduction

Diatoms, as one of the most diverse groups of phytoplankton, contribute approximately 20-40% to oceanic primary production and play a crucial role in sustaining marine food webs (Nelson et al. 1995, Mann 1999). The sinking of diatoms influences the community composition of phytoplankton by affecting their residence time in the photic zone and serves as an important pathway for material loss from the surface to the deep ocean (Cassar et al. 2015, Le Moigne et al. 2015). A key characteristic of diatoms is their silicified cell wall, which is known as frustules and contributes to their faster sinking rate compared to many other types of phytoplankton (Ducklow et al. 2001). Given their high production and faster sinking rate, diatoms account for a significant proportion of carbon and silicate export, making diatom sinking crucial for the global cycling of silica and carbon (Ragueneau et al. 2006).

Most pelagic diatoms are considered passive sinkers due to their lack of motor organelles, such as flagella, as observed in dinoflagellates (Smayda 1970a, Waite et al. 1997). The speed of passive sinking was believed to depend on cell size, as described by Navier-Stoke’s equation (Bienfang 1981a). However, recent studies have shown that large diatoms indeed can actively regulate their sinking rates, and this capability is not related to their cell size. (Gemmell et al. 2016, Du Clos et al. 2019, Du Clos et al. 2021). This regulated sinking dynamic, also known as sinking behavior, is an energy-intensive process involving various physiological responses (Waite et al. 1992, Du Clos et al. 2019). These responses probably include the production of organic osmolytes, ionic regulation, carbohydrate ballasting, solute adjustment, active water transport and periodic cell expansion facilitated by cytoskeletal motors (Boyd & Gradmann 2002, Woods & Villareal 2008, Raven & Doblin 2014, Lavoie et al. 2016, Lavoie & Raven 2020).

Sinking regulation allows diatoms to control their position in specific layers of the water column, optimizing their access to light and nutrients, which are crucial for their survival but vary inversely with depth (Smayda 1970b, Dubinsky & Schofield 2010, Du Clos et al. 2019). Moreover, the unique sinking behavior resulting from sinking regulation in diatoms allows for efficient nutrient flux to their cell surfaces (Gemmell et al. 2016). Consequently, sinking regulation empowers diatoms to occupy new ecological niches, promoting their remarkable diversity.

Similar to nitrogen and phosphorus, silicon is a crucial nutrient for diatom growth. As a single-celled organism, the diatoms possess the ability to perceive and respond to changes in their surrounding environment with great sensitivity, allowing them to regulate their sinking behavior accordingly (Karp-Boss et al. 1996, Falciatore et al. 2000, Du Clos et al. 2021). In the case of the effects of silicate deficiency on the sinking of diatoms in the time domain, a short-term deprivation of silicate (2 h) results in a decrease in the mean sinking speed and an increase in fluctuations in the instantaneous sinking speed (Du Clos et al. 2021). However, a prolonged silicate deficiency for 24-48 hours results in an increase in the mean sinking speed of diatoms (Bienfang et al. 1982).

Nevertheless, the characteristics of sinking behavior in the frequency domain and their response to nutrient limitations have been largely overlooked in studies on diatom sinking behavior (Gemmell et al. 2016, Du Clos et al. 2019, Du Clos et al. 2021). Understanding sinking behavior solely in the time domain falls short of providing a comprehensive picture, because it lacks crucial information such as frequency and period. In fact, the frequency domain is a vital perspective in studying biological behavior and has been extensively utilized in various fields, including human behavior (Watson & Ahumada Jr 1983).

Wavelet analysis is powerful in decomposing signals into subsignals of different frequency ranges, providing a robust tool for analysing the time and frequency characteristics of the signal (Torrence & Compo 1998). Compared to Fourier analysis, wavelet analysis exhibits better time-frequency localization properties, allowing for more precise capture of instantaneous features and frequency variations. Moreover, wavelet analysis can effectively handle nonstationary signals, making it a valuable tool for studying nonstationary signals and nonlinear systems (Labat 2005). The advantages of multiresolution and adaptivity in wavelet analysis have led to its wide application in the analysis of time series data in fields such as hydrology, planktonology and marine ecology (Kang & Lin 2007, Wang et al. 2012, Cazelles et al. 2014, Carey et al. 2016). The sinking dynamics of large diatoms, represented as a time-varying curve of instantaneous sinking speed, are nonstationary time series data (Gemmell et al. 2016, Du Clos et al. 2019, Du Clos et al. 2021). Therefore, it can be analyzed using wavelet analysis, which can identify the presence of active regulation (significant oscillations in the frequency domain) during sinking and localize it in time .

*Palmerina hardmaniana,* a large diatom widely distributed in tropical and subtropical coastal waters (Garcia & Odebrecht 2008), has been confirmed to possess the ability to regulate its sinking behavior (Gemmell et al. 2016). This study examines how *P. hardmaniana* regulates its sinking behavior under silicate depletion, by exploring the temporal and frequency aspects of this regulation. To study this phenomenon, the sinking process of *P. hardmaniana* was captured and converted into digital signals using image processing techniques. Wavelet analysis was then applied to analyze the time-frequency characteristics of the sinking dynamics to identify the active regulations during sinking and track changes in these regulations over time. This study is the first report analysing the characteristics of large diatom sinking behavior in the time-frequency domain, and it may contribute to a deeper understanding of diatom sinking behavior.

## 2. Material and methods

### 2.1 Diatom cultures

The *P. hardmaniana* used in this study was obtained from the Culture Collection of Marine Algae at the Guangxi Academy of Science. The diatom was cultured in f/2 culture medium with a light-dark cycle of 12 hours light and 12 hours dark, providing an irradiance of 100 µmol photons m^-2^ s^-1^. The culture was maintained at a temperature of 20L and a salinity of 30 psu. Only cells in the exponential growth phase were chosen for subsequent experiments.

### 2.2 Experimental setup

*P. hardmaniana* cells in the exponential growth phase were collected by filtering culture medium through a 20 µm mesh. The collected cells were then washed three times with nutrient-free artificial seawater to remove nutrients attached to the diatom surface. To compare the sinking behavior of the diatom under silicate-enriched and silicate-depleted conditions, the cleaned diatom cells were introduced into f/2 medium and silicate-depleted f/2 medium, respectively. The f/2 medium is a eutrophic culture medium (Guillard 1975), while the silicate-depleted f/2 medium contains all nutrients except for silicate. The cells were introduced at an initial density of 17 cells ml^-1^. The experiments were conducted in triplicate using 500 ml plastic culture bottles. All culture conditions, except for the silicate concentration, remained the same as the strain maintenance conditions mentioned above. Sampling was performed at approximately 11:00 am, four hours after the start of the light phase. Daily samples were taken to measure the cell density and F_v_/F_m_ value. The approximate growth status of *P. hardmaniana* in the control group could be determined based on the results of daily cell counting. Based on the cell growth status of the control group, sinking observations were made during the adaptation period (day 0 and day 3), the early stage of the exponential growth phase (day 6), and the full-fledged exponential growth phase (day 9). The sampling water was gently introduced into a convection-free system (observation chamber) for observation, following the methods described below. Notably, sampling on day 0 referred to the sampling taken 30 minutes after the start of the experiments.

### 2.3 Creation of convection-free water column

The sinking behavior of *P. hardmaniana* was monitored in a transparent plastic chamber measuring 10 cm in length, 6 cm in width and 20 cm in height. The experimental temperature was maintained consistent with the culture temperature. When water is added to the chamber, convection currents are generated, which can significantly affect the sinking of diatoms due to their small cell size. To minimize the influence of convection currents, a convection-free water column with stabilized salinity was established following the method described by O’Brien et al. (2006). In summary, two 500 ml vessels were filled with high-salinity water (32 psu) and low-salinity water (28 psu). These vessels were connected at the bottom by a plastic pipe and controlled by a valve. Throughout the process, the water in the vessels was continuously stirred using a magnetic stirrer. At the beginning of the experiments, the valve was opened, and the low-salinity water was slowly pumped into the observation chamber. This pumping caused the high-salinity water to enter the low-salinity chamber from the bottom and mix with the low-salinity water. As a result, a linear salinity gradient was established in the observation chamber, creating a convection-free water column that allowed for the vertical sinking of diatoms. The schematic diagram of the experiment can also be found in the description provided by Du Clos et al. (2021).

### 2.4 Observation of *P. hardmaniana* sinking behavior

The sinking process of *P. hardmaniana* was recorded using the built-in camera of a smartphone (HONOR V20). The camera was set to professional mode, capturing videos at 60 frames per second with a pixel resolution of 1920×1080. To ensure optimal illumination, an LED illuminator was utilized during photography. Prior to observation, three samples from replicate treatments were combined, and the mixed sample was gently added to the water surface of the observation chamber using a spoon. This addition resulted in convection currents occurring from the surface to a depth of 4-8 cm. Therefore, the focal plane was positioned below a depth of 10 cm to capture the sinking behavior. Sinking tracks were extracted from the recorded videos, and the sinking speed of diatoms was analyzed using the ImageJ package Trackmate (Tinevez J Y et al. 2017). Only diatoms that were in clear focus and away from the container walls were selected for analysis. A total of 20 cells with a sinking time of 150 seconds were chosen for sinking analysis. The mean sinking speed was calculated as the average of the 20 cells within a 150-second timeframe.

For the recorded video, a sequence of images was extracted from PotPlayer at a frequency of 1 image per second. These images were imported into the ImageJ package TrackMate for trajectory tracking of the focused cells (Fig.1). Prior to tracking, the images were denoised. To ensure reliable trajectories, images with complete and continuous cell trajectories were screened. Trajectories perpendicular to the horizontal plane and appearing straight were selected, indicating unaffected cells by convection and disturbances. The instantaneous sinking rate of cells was calculated by measuring the distance traveled per second. The average sinking rate mentioned in this paper represents the mean value derived from all instantaneous sinking rate observed in the selected cell trajectories.

**Figure 1.**
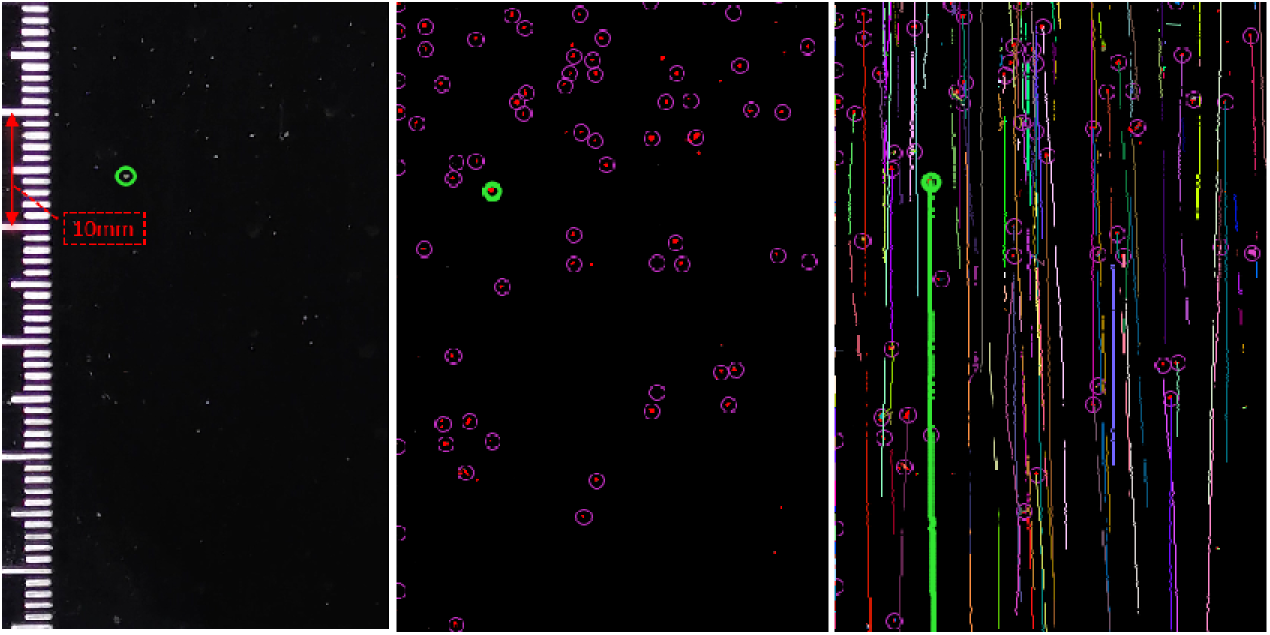
illustrates the trajectory tracking process using the ImageJ package TrackMate interface. The individual cell being tracked is enclosed within a green circular bounding box, while its complete motion trajectory is represented by the thick green line in the image.

### 2.5 Cell density and F_v_/F_m_ value

A 3 ml sample was taken daily to measure the cell density and F_v_/F_m_ value of *P. hardmaniana*. The cell number was determined using an inverted microscope (Nikon, TE 300). The maximum quantum yield of PSII, F_v_/F_m_, was measured using a Phyto-PAM fluorometer (Heinz Walz GmbH, Effeltrich, Germany). Before measurement, the samples were dark-adapted for 15 minutes. F_v_/F_m_ was calculated as (F_m_-F_o_)/F_m_, where F_m_ represents the maximal fluorescence and F_o_ represents the minimal fluorescence from a dark-adapted sample. To determine F_o_, a low-intensity modulated measuring light of 28 μmol m^-2^ s^-1^ was used. F_m_ was induced by a red saturating light pulse (10000 μmol m^-2^s^-1^) after exposure to an actinic light of 166 μmol m^-2^ s^-1^.

### 2.6 Wavelet analysis

Wavelet analysis is a powerful tool for analysing time series data that allows for the examination of frequency characteristics over time. The variation in the instantaneous sinking speed of *P. hardmaniana*, being a nonstationary time series in the time domain, was analyzed using continuous wavelet transform (CWT). Morlet wavelet is used here as the base function was based on its ability to accurately represent the shape of sinking variation and provide a favourable balance between time and frequency localization. All wavelet calculations referenced the description provided by Grinsted et al. (2004). The wavelet power spectrum was calculated to reveal the spectral power patterns of sinking behavior in both the time and frequency domains. Wavelet coherence and wavelet phase difference were used to assess the similarity and phase differences in the sinking pattern of *P. hardmaniana* between the control and silicate-depleted treatment at various frequencies and time scales. For all wavelet calculations discussed in this post, we utilized the Matlab wavelet packet described by Grinsted et al. (2004), which can be freely accessed at http://grinsted.github.io/wavelet-coherence/ (last accessed: 30th July 2018).

### 2.7 Statistical analysis

To assess the differences in mean sinking speed, F_v_/F_m_ value, and cell density between the control and silicate-depleted treatment on the same day, a t test was conducted with a significance level of 5%.

## 3. Results

### 3.1 Cell density and F_v_/F_m_ value

Growth of *P. hardmaniana* was found to be inhibited under silicate-depleted treatments, resulting in a significant decrease in the cell density starting from day 4 (*p*<0.05, t test). The density of *P. hardmaniana* decreased from an initial density of 17.00±2.08 cells ml^-1^ to 2.00±1.73 cells ml^-1^ at the end of the experiment. In contrast, the cells in the control group exhibited exponential growth, with the density reaching 108.00±8.89 cells ml^-1^ at the end of the experiment (Fig. 2 a). The growth curve in the control group showed an initial adaptation phase from day 0 to day 3, followed by an early stage of exponential growth from day 4 to day 6, and a full-fledged exponential growth phase from day 7 to day 9 (Fig. 2 a). The F_v_/F_m_ value also indicated significant environmental stress due to silicate depletion, which was first observed on day 2 (*p*<0.05, t test). The variations in the F_v_/F_m_ value followed a similar pattern to that of cell density, but the response to silicate depletion appeared earlier in the F_v_/F_m_ value compared to the density (Fig. 2 b).

**Figure 2.**
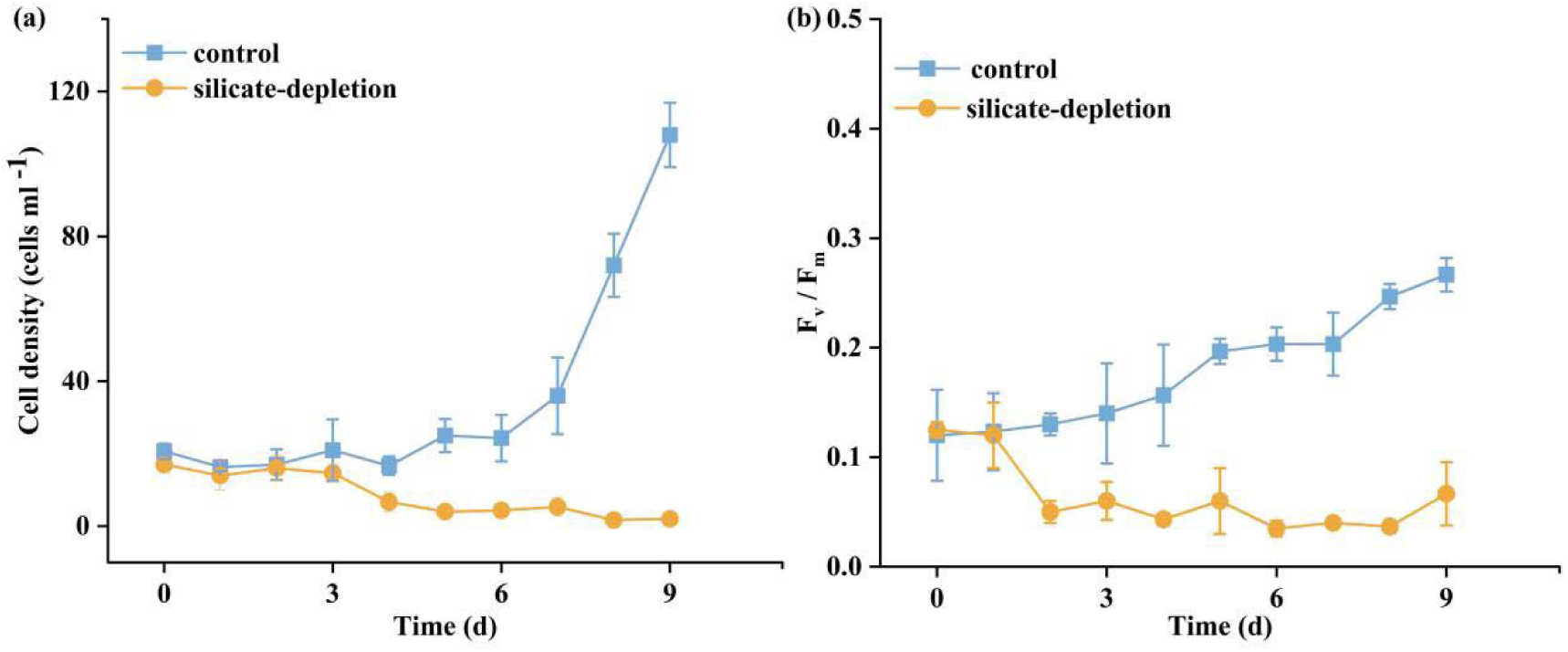
Variations in cell density (a) and F_v_/F_m_ value (b) of *P. hardmaniana* under conditions of nutrient repletion and silicate depletion

### 3.2 Sinking dynamics and mean sinking speed of *P. hardmaniana*

The instantaneous sinking speed of *P. hardmaniana* displayed oscillations over time (Fig. 3). Aside from the mean sinking speed, it was challenging to discern differences in sinking dynamics (e.g., variations in instantaneous sinking speed) between silicate depletion and the control in the time domain (Fig. 3 and Fig. 4). Under silicate depletion, the mean sinking speeds on day 0, day 6 and day 9 were 0.11±0.03 mm s^-1^, 0.14±0.04 mm s^-1^ and 0.13±0.03 mm s^-1^, respectively, which were significantly lower than those of the control group (*p*<0.01, t test, Fig. 4). However, the mean sinking speed of *P. hardmaniana* in the silicate-depleted condition was 0.19±0.07 mm s^-1^ on day 3, which was significantly higher than that of the control group (*p*<0.01, t test, Fig. 4). The mean sinking speed of diatom cells in the control group exhibited variations across different growth stages. These differences in mean sinking speed between the control and silicate-depleted groups on day 3, day 6, and day 9 were not solely attributed to silicate depletion but were also attributed to the growth status of cells. Nonetheless, the observed differences were primarily driven by silicate depletion, particularly on day 0, day 3 and day 9 (Fig. 4).

**Figure 3.**
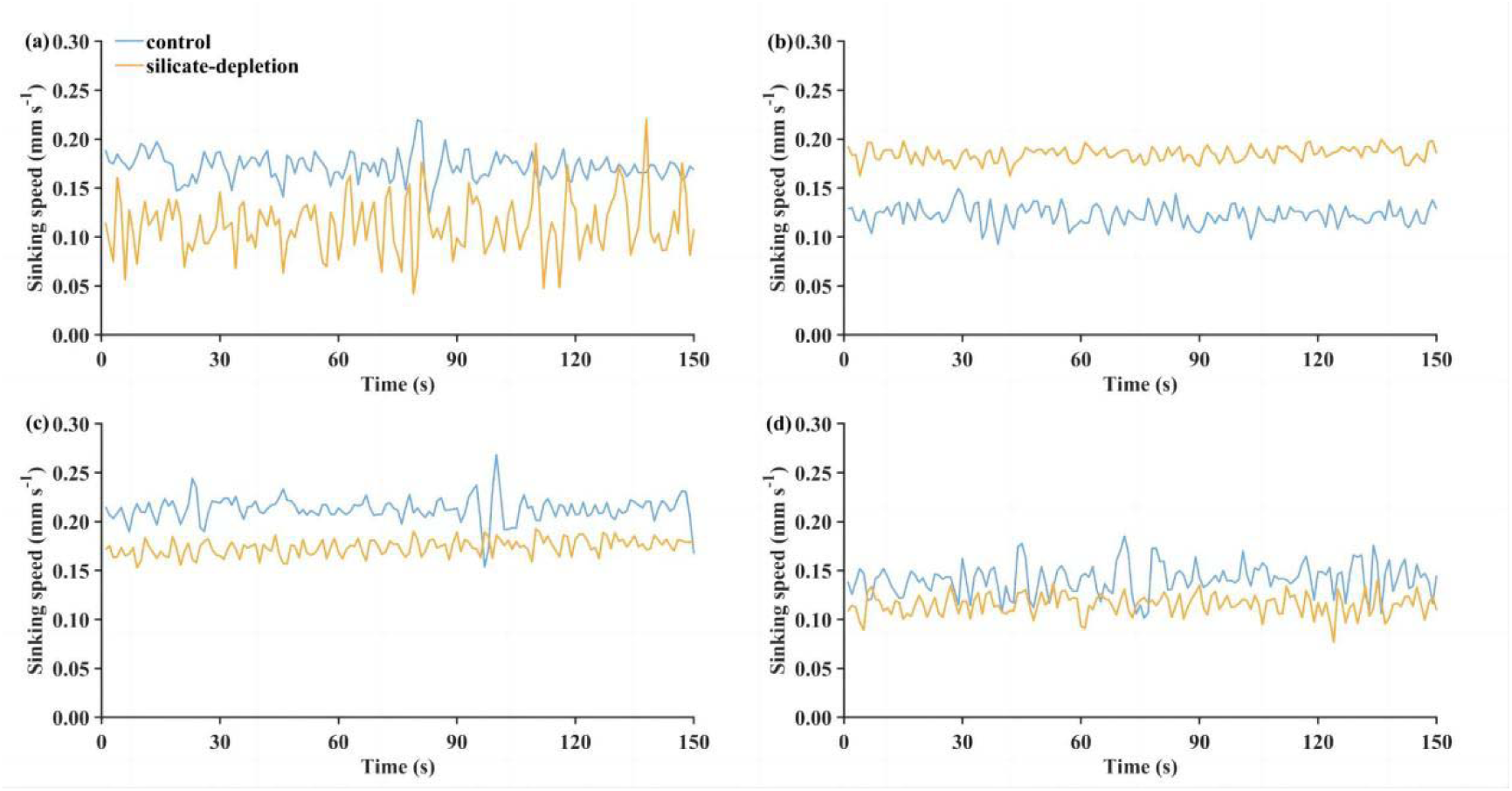
Instantaneous sinking speeds of representative *P. hardmaniana* cells on day 0 (a), day 3 (b), day 6 (c) and day 9 (d).

**Figure 4.**
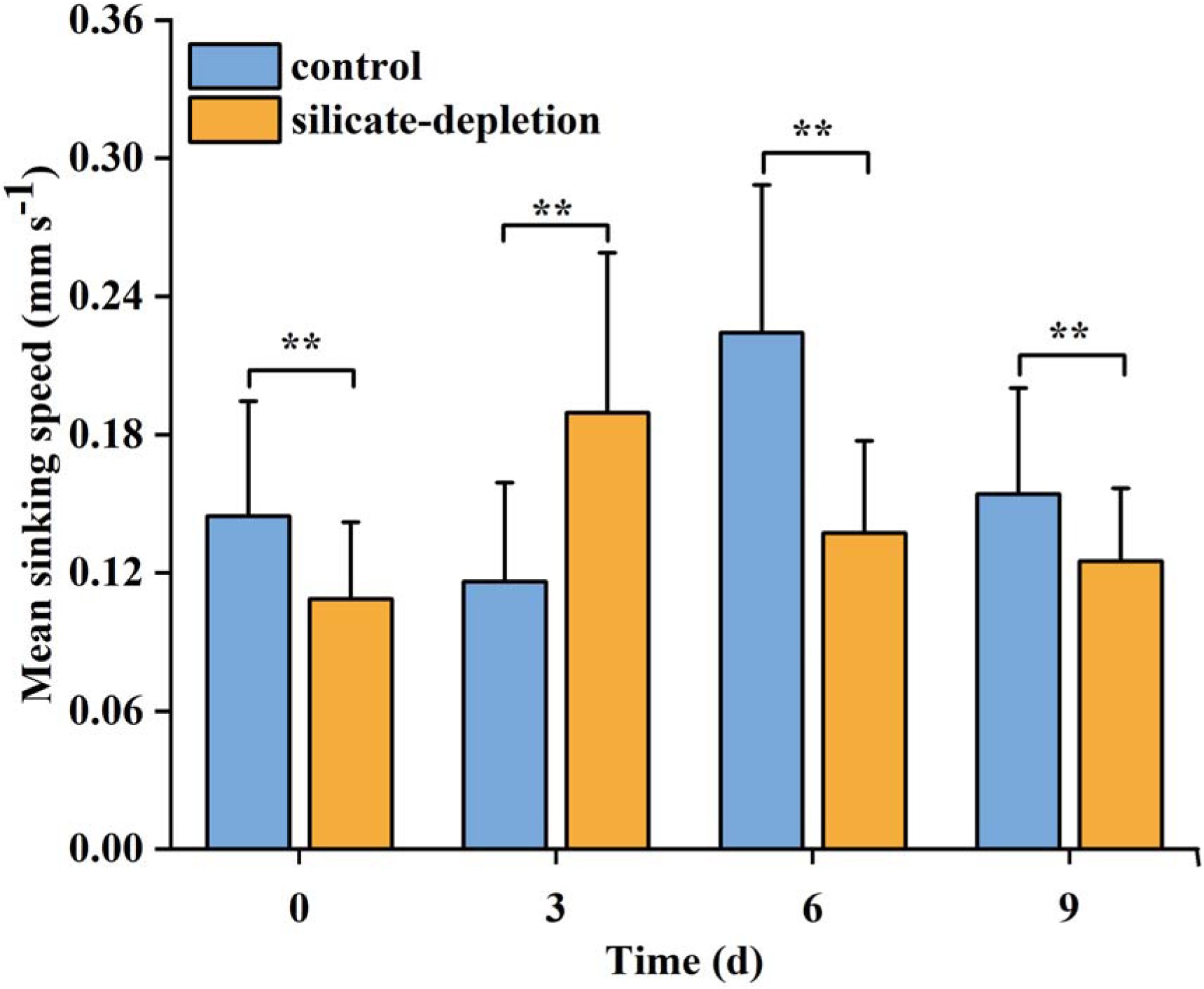
Mean sinking speed of *P. hardmaniana*. The two asterisks above the column indicate highly significant differences between the control and silicate-depleted treatments on the same day (*p*<0.01).

### 3.3 Distribution of the instantaneous sinking speed of *P. hardmaniana*

*P. hardmaniana* exhibited the ability to regulate its sinking speed by adjusting the occurrence of different magnitudes of instantaneous sinking speeds (Fig. 5). Compared to the control group, under silicate-depleted treatments, *P. hardmaniana* decreased the occurrence of instantaneous sinking speed in the range of 0.15-0.20 mm s^-1^ while increasing the occurrence in the range of 0.08-0.14 mm s^-1^, resulting in a reduction in mean sinking speed on day 0 (Fig. 5 a). The increase in mean sinking speed on day 3 was achieved by increasing the occurrence of sinking speeds in the range of 0.13-0.40 mm s^-1^ while decreasing the occurrence in the range of 0.03-0.12 mm s^-1^ (Fig. 5 b). Similarly, under silicate-depleted conditions, *P. hardmaniana* reduced its mean sinking speed on day 6 by decreasing the sinking speed in the range of 0.19-0.36 mm s^-1^, while increasing the sinking speed in the range of 0.04-0.18 mm s^-1^ (Fig. 5 c). On day 9, *P. hardmaniana* in silicate-depleted conditions decreased its mean sinking speed by decreasing the occurrence of sinking speeds in the range of 0.16-0.24 mm s^-1^, while increasing the occurrence in the range of 0.09-0.14 mm s^-1^ (Fig. 5 d).

**Figure 5.**
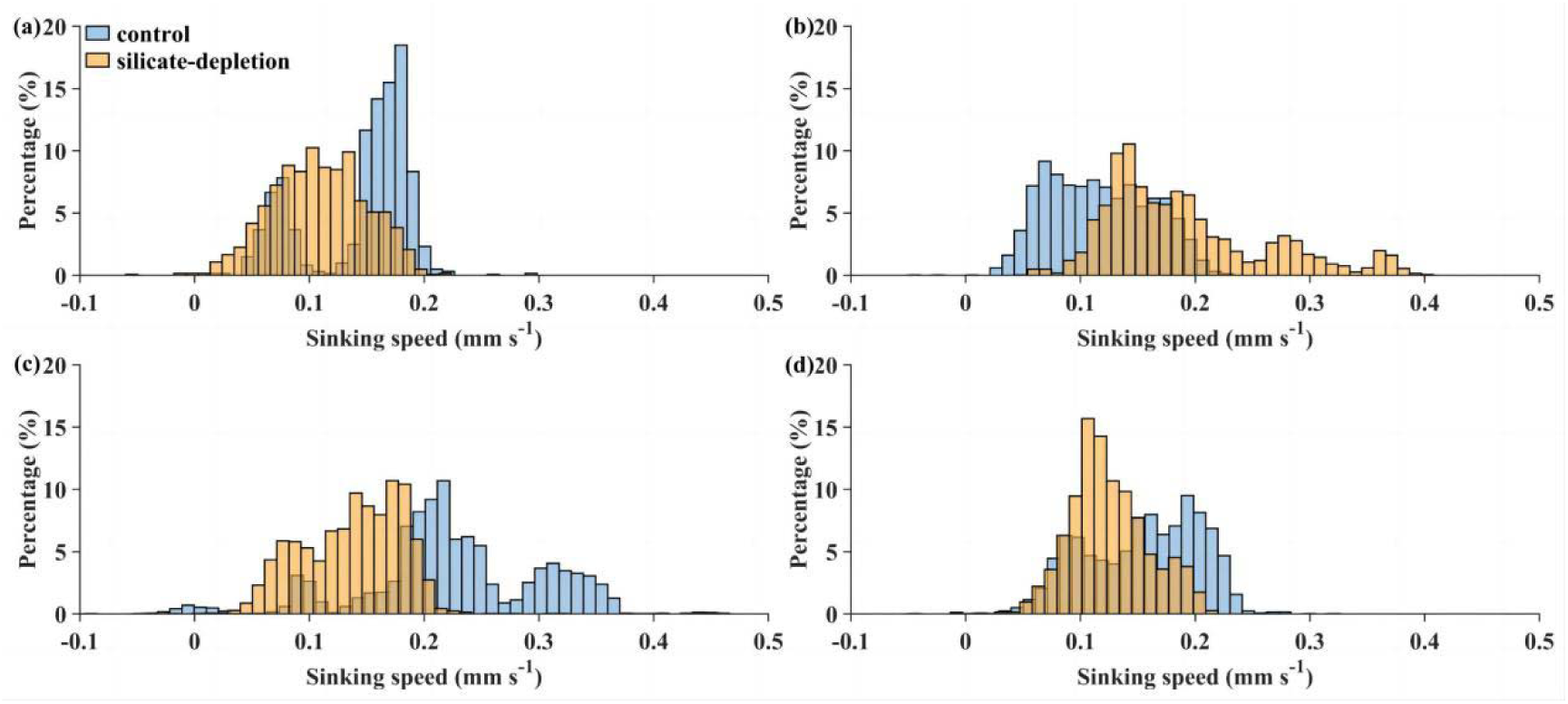
Distribution of the instantaneous sinking speed of *P. hardmaniana* over 150 seconds on day 0 (a), day 3 (b), day 6 (c) and day 9 (d).

### 3.4 Wavelet power spectrum and wavelet coherence analysis

The active regulation of sinking speed in *P. hardmaniana* was clearly evident as a pronounced increase in power in the wavelet power spectrum. Throughout the time series, no continuous dominant scales were observed in either the control or silicate-depleted treatment, indicating that the regulation of sinking speed in *P. hardmaniana* exhibited intermittent patterns in the frequency domain (Fig. 6). The sinking regulation frequency of P. hardmaniana exhibited variability in response to cultivation conditions and growth stages, typically ranging from approximately 0.13-0.50 Hz (corresponding to a period of 2-8 seconds). However, a distinct regulation frequency of 0.10-0.14 Hz (resulting in a period of 7-10 seconds) was uniquely observed on day 9, specifically between the 63rd and 90th second (Fig. 6). Although the timing of active regulation varied, the sinking speed of *P. hardmaniana* in the control group exhibited only one dominant scale on day 0 (0.13-0.25 Hz), day 3 (0.15-0.25 Hz) and day 6 (0.13-0.25 Hz), respectively. Compared to the control group, silicate-depleted conditions showed more dominant or subdominant scales in the sinking speed, indicating more sinking regulation during the study periods (Fig. 6 e-h). The number of sinking regulations in the control group increased on day 9, while the number of sinking regulations in the silicate-depleted group decreased on day 9. Additionally, the wavelet power spectrum revealed that the power of the signal at dominant scales in the control group was higher than that in the silicate-depleted group on day 3, day 6 and day 9. Conversely, the power of the signal at dominant scales in the silicate-depleted group was higher than that in the control group on day 0 (Fig. 6).

**Figure 6.**
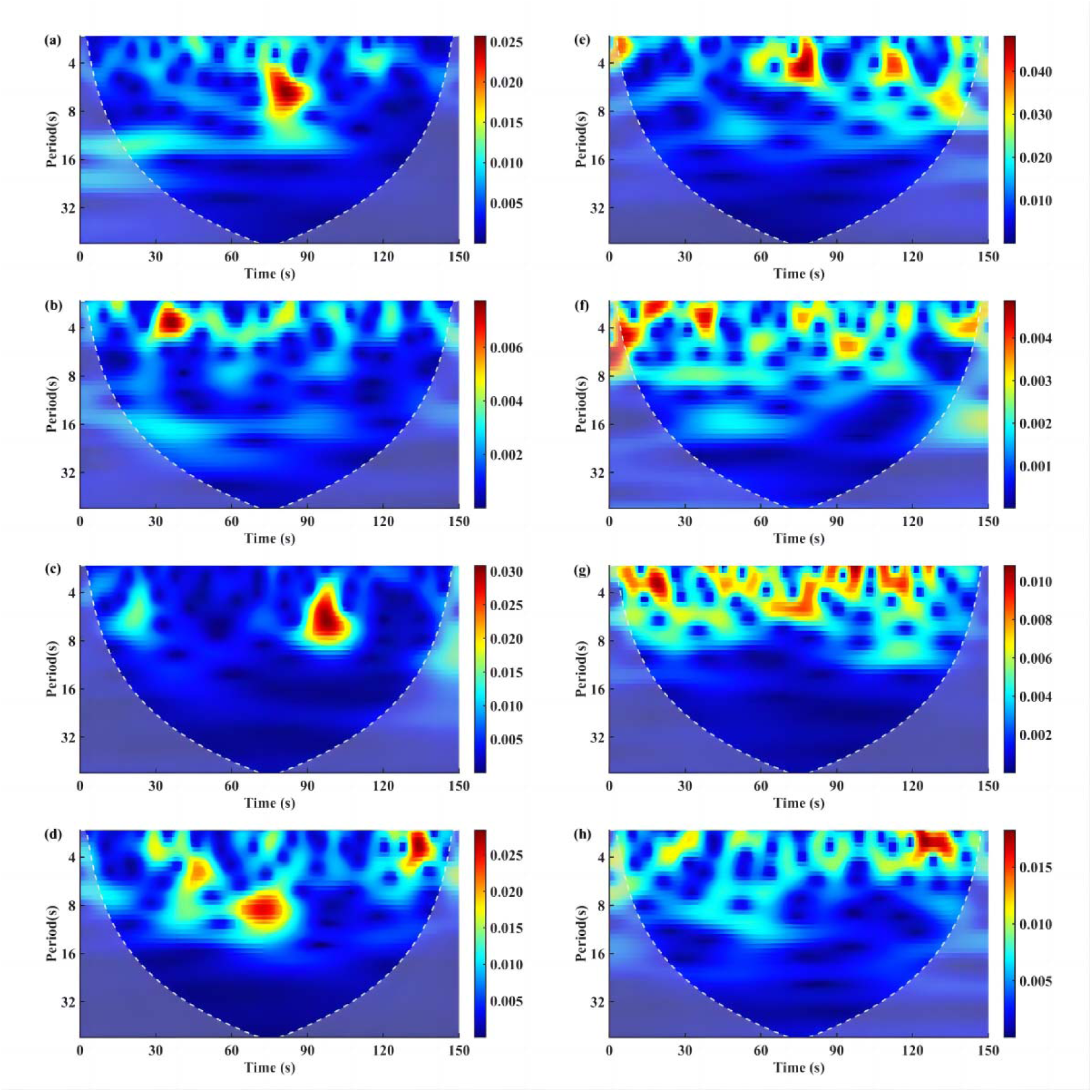
Wavelet power spectrum of the sinking speed of *P. hardmaniana.* The lowercase letters a, b, c and d represent the silicate-depleted treatments on days 0, 3, 6 and 9, respectively. The lowercase e, f, g and h represent the control on day 0, day 3, day 6 and day 9, respectively. The area outside the dashed line represents the cone of influence (COI), where edge effects become significant and may affect the accuracy of signal characteristics. Note that the power axis scales of each figure are different, and deeper colors in the same figure indicate relatively higher power values.

The dominant scale in both the control and silicate-depleted treatments mainly fell within the range of approximately 2-8 seconds (Fig. 6). Therefore, this analysis focused on the coherence of the bands within this 2-8 second range. The phase analysis revealed that the majority of associations in the variation of sinking speed between the silicate-depletion and control were out of phase or in anti-phase, with the exceptions of small bands that appeared on day 3 and day 9 (Fig. 7). The exception on day 3 was the 2-6 second band between the 34^th^ and 41^st^ seconds, which was determined to be in phase with a phase advancement of approximately 42^°^ (Fig. 7 b). The exceptions on day 9 were the 2-8 second bands between the 25^th^ and 30^th^ seconds and between the 115^th^ and 123^rd^ seconds, which were also observed to be in phase (Fig. 7 d). However, the impact of these exception bands was weak as their duration did not exceed 8 seconds during a 150 -s observation period.

**Figure 7.**
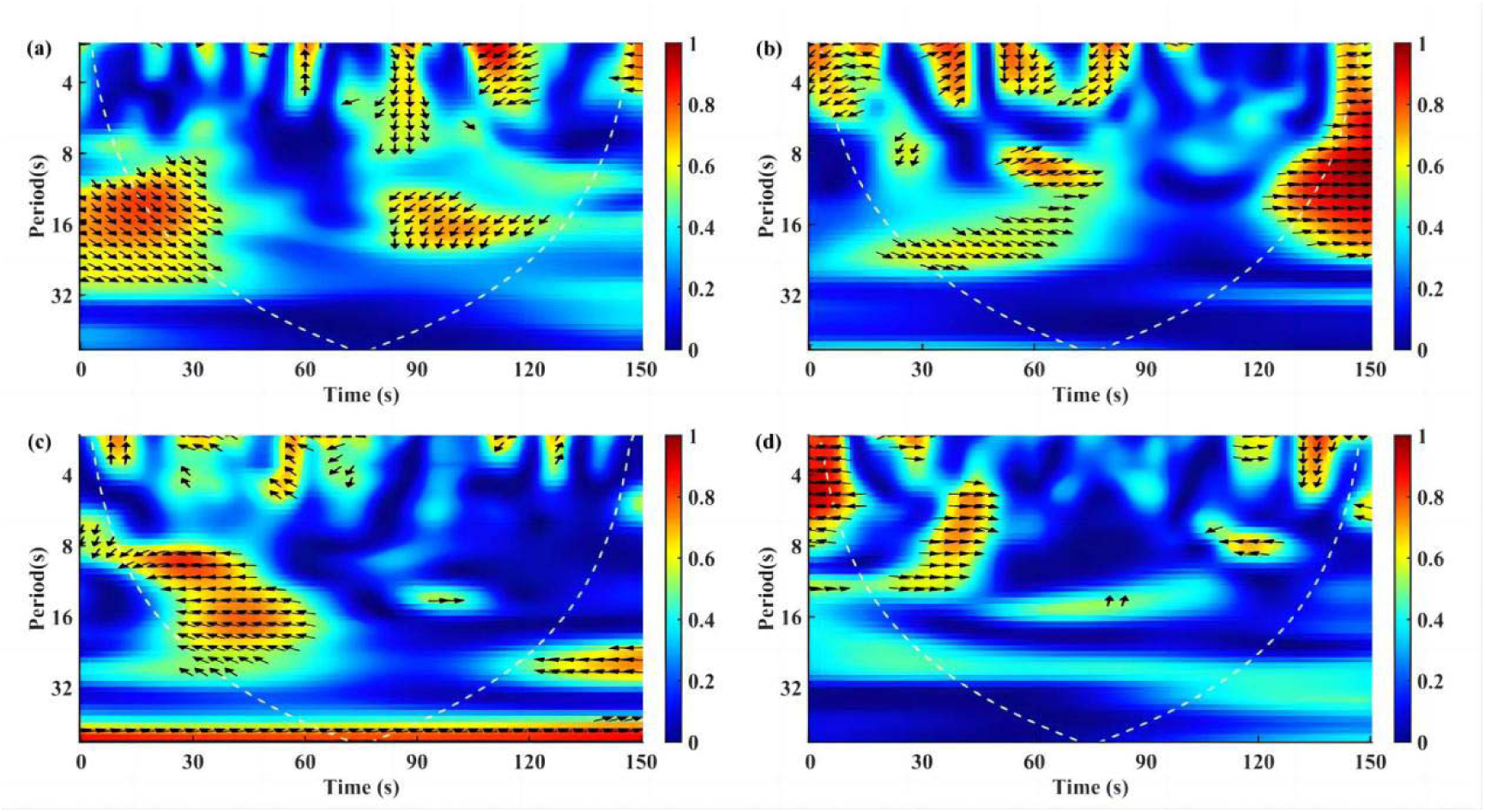
Wavelet coherence of sinking speed of *P. hardmaniana* between silicate depletion and control on day 0 (a), day 3(b), day 6(c) and day 9 (d). Deeper colours indicate higher coherence. The arrow pointing to the horizontal right (0^°^) indicates an in-phase situation, while pointing to the horizontal left (180°) indicates an anti-phase situation. The arrow pointing to the vertical up (90°) or down (270°or -90°) indicates an out of phase situation.

## 4. Discussion

### 4.1 Variations in the instantaneous sinking speed of *P. hardmaniana*

Some species of large diatoms actively regulate their sinking speed in response to environmental variations (Du Clos et al. 2021). These diatoms possess the ability of autonomous regulation, resulting in a rapid change in instantaneous sinking speed (Gemmell et al. 2016, Du Clos et al. 2019, Du Clos et al. 2021). Consistent with the observation of Gemmell et al. (2016), a capability for active regulation of sinking speed was also found in *P. hardmaniana* (Fig. 3). The range of variations in instantaneous sinking speed observed in our study was similar (in the control group on day 3 and day 9 and in the silicate-depletion group on day 9) or greater (in the control group at day 0 and day 6 and in the silicate-depletion group on day 0, day 3 and day 6) than that reported in a previous study (0.02-0.15 mm s^-1^). However, the variations in the instantaneous sinking speed in our study displayed higher oscillation than that of the study by Gemmell et al. (2016) for *P. hardmaniana*. Rather, our study showed more similarity to the sinking dynamics of *Coscinodiscus radiatus* as described in that of Gemmell et al. Notably, the *P. hardmaniana* used in their study was collected from the field, whereas *P. hardmaniana* in our study has been maintained under artificial conditions for over two years. Although further research is needed, to determine the exact reasons, the differences in the sinking behavior of *P. hardmaniana* between field and indoor cultures suggest that the sinking behavior of large diatoms is not species-specific but rather influenced by their survival environments. Additionally, the regulation of sinking behavior in large diatoms is dependent on energy, and lowering the oscillation may conserve energy (Waite et al. 1992, Lavoie et al. 2016, Lavoie & Raven 2020). Energy is precious for survival in challenging field conditions, and it is possible that *P. hardmaniana* under harsher field conditions has more energy-efficient ways of regulating sinking behavior. For example, they may achieve sinking regulation through low-frequency oscillations by leveraging water turbulence in the field (Karp-Boss et al. 1996, Wolf-Gladrow & Riebesell 1997).

### 4.2 Response of the mean sinking speed of *P. hardmaniana* to silicate depletion

Diatoms with excess silicon become more heavily silicified, which leads to faster sinking of their dead cells (Durkin et al. 2013). However, it is counterintuitive that some species of live diatoms also increase their mean sinking speed under silicate limitation (Bienfang et al. 1982, Bienfang & Harrison 1984a, Aumack & Juhl 2015). One possible explanation for this dynamic is that the diatom cells under silicate deficiency are arrested in the G2 phase (Brzezinski et al. 1990, Martin-Jézéquel et al. 2000, Aumack & Juhl 2015). During the G2 phase, cells successfully completed DNA replication and synthesized most cellular components. However, their ability to undergo further division is hindered by the limited access to silicon, which is necessary for the production of new silica frustules. As a result, cells become heavier and their sinking increases (Bienfang & Harrison 1984b, Thompson et al. 1991, Aumack & Juhl 2015). It is possible that *P. hardmaniana* cells in the silicate-depleted treatment were arrested in the G2 phase on day 3 (Fig. 2 a). This state could increase the mean sinking speed of diatoms (Aumack & Juhl 2015). Consistent with the experimental results from Aumack & Juhl’s study on *Nitzschia frigida*, the mean sinking speed of *P. hardmaniana* significantly increased on day 3 (Fig. 4). This suggests that under silicate limitation, different species of diatoms may exhibit similar physiological responses, where their sinking rates increase when silicate is scarce. Such a response could be a strategy adopted by diatoms to adapt to silicate limitation, promoting the transport of organic matter to the seafloor by increasing sinking rates, which may have significant implications for nutrient cycling and energy flow within marine ecosystems.

As mentioned above, most studies have found that silicon limitation increases the mean sinking speed of diatoms (Bienfang et al. 1982, Bienfang & Harrison 1984a, Aumack & Juhl 2015). However, these studies were typically conducted after 24-48 hours of silicate stress. The effects of prolonged silicate stress on the mean sinking speed of diatoms remain unclear. Our results indicated a significant decrease in the mean sinking speed of *P. hardmaniana* on day 6 and day 9 (Fig. 4). As a compensatory response to silicon deficiency, diatoms arrested in G2 phase shift their photophysiology to increase their demand for light energy in the late G2 phase, which is linked to the energetic requirements for enhancing silicon transport and deposition before cell division (Smith et al. 2016). The decrease in the mean sinking speed of *P. hardmaniana* on day 6 and day 9 under silicate depletion (Fig. 5) may facilitate its longer residence in the upper water column. This extended time in the upper water column allows *P. hardmaniana* to acquire more light energy for later light-dependent cell cycle processes. Silicon starvation also induces lipid accumulation in diatoms (Schnurr et al. 2013, Smith et al. 2016). Due to the lower density of lipids compared to seawater, the accumulation of lipids in diatoms may also reduce their sinking speed.

In fact, the results on day 0 were measured after a 30 min exposure to silicate depletion. Large diatoms are capable of rapidly altering their sinking in response to changes in nutrient availability, and the short-term response to nutrient variations differs from the long-term response. For example, the mean sinking speed of *Coscinodiscus wailesii* increases within the first 2 hours but declines over the next 22 hours when they are cultured from silicate-depleted to eutrophic conditions (Du Clos et al. 2021). Contrary to their treatments, *P. hardmaniana* in this study was inoculated from eutrophic (f/2) to silicate-depleted conditions; interestingly, the corresponding regulation of sinking was also opposite, as the mean sinking speed of *P. hardmaniana* decreased on day 0 but increased on day 3 (Fig. 4). In fact, the sinking of diatoms is affected by both external nutrient concentrations and internal nutrient pools (Bienfang 1981b, Bienfang & Szyper 1982). As a sensitive indicator of environmental stress (Huang et al. 2016), the F_v_/F_m_ value indicated that *P. hardmaniana* was not stressed by silicate depletion on day 0 (Fig. 2). Most likely, *P. hardmaniana* has the capacity to store a substantial amount of silicon in internal nutrient pools due to its large size. Lower organisms possess chemoreceptors on their membranes that enable them to effectively sense and respond to environmental variations (Maddock & Shapiro 1993). After being introduced to a new environment, *P. hardmaniana* may promptly sense a lack of silicate. In response, *P. hardmaniana* may reduced its sinking speed on day 0, enabling it to better utilize solar energy, which is necessary to power diatom replication, synthesis and sinking regulation. To address silicate depletion, *P. hardmaniana* may exhibit a preference for maximizing energy acquisition under conditions of sufficient silicon reserves. This response is beneficial for subsequent processes such as DNA replication, cell synthesis and sinking regulation.

### 4.3 The response of *P. hardmaniana* sinking behavior to silicate depletion in the time and frequency domains

To date, the sinking behavior of large diatoms has been well documented in the time domain, while their behavior characteristics in the frequency domain remain unexplored (Gemmell et al. 2016, Du Clos et al. 2019, Du Clos et al. 2021). Due to the complexity of the sinking dynamics (Fig. 3), identifying the differences in sinking behavior between the control and silicate-depleted treatments only in the time domain, apart from their average and oscillation, is challenging (Figs. 3 and 4). The analysis of behavior in the frequency domain is helpful in understanding intricate organism behavior because it allows for the removal of noise interference and acquisition of key information about motion (Watson & Ahumada Jr 1983, Adelson & Bergen 1985, Jhuang et al. 2007). In this study, wavelet analysis conducted in the time-frequency domain revealed that the sinking regulation of *P. hardmaniana* was intermittent (Fig. 6). The main difference between the control group and silicate-depleted treatment was that *P. hardmaniana* under silicate depletion enhanced its sinking regulation (0.13 Hz ∼ 0.50 Hz) during the sinking process (Fig. 6). The enhancements of the regulation of course increase the oscillation of the instantaneous speed. In addition, compared to the control group, the increasing regulation in the silicate-depleted group was primarily driven by silicate deficiency (Fig. 7). These findings further confirm from the perspective of the time-frequency domain that nutrient limitation enhances the sinking oscillation of diatoms (Du Clos et al. 2019).

Additionally, the occurrence of diatom sinking regulation appears to be related to the physiological state of the cells in both the control and silicate-depleted conditions (Fig. 6). In the silicate-depleted treatment, the capacity of *P. hardmaniana* to enhance its active regulation was limited on day 9 compared to that on day 0, day 3 and day 6 (Fig. 6). This is probably due to the deteriorated physiological state of cells on day 9 resulting from prolonged silicate depletion stress (Fig. 2 b). Although silicate deficiency increased the occurrence of sinking regulation, the oscillation power was higher in the control group than in the silicate-depleted treatment group, possibly due to better physiological conditions of cells in the control group, except on day 0 (Fig. 2 and Fig. 6). On day 0, after 30 minutes of silicate stress, *P. hardmaniana* exhibited an increase not only in the occurrence of regulation but also in the oscillation power, indicating the capability of large diatoms to promptly respond to environmental stress within minutes when adequate energy and resource reserves are present. In the control group, *P. hardmaniana* regulated its sinking only once (0.13-0.50 Hz) during the 150-second observation period before the full-fledged exponential growth phase (day 0, day 3 and day 6, Fig. 6 a, b and c). However, in the full-fledged exponential growth phase (day 9), an additional regulation with a frequency of 0.08-0.14 Hz (equivalent to a period of 7-12 s) was observed in addition to the 0.13-0.50 Hz regulation (Fig. 6 d). Although nutrient levels were not measured during the experiment, it is expected that they naturally decrease as cells proliferate. The absence of this additional regulation (0.08-0.14 Hz) in the silicate-depleted treatment suggests that it is unlikely to be a response to silicate deficiency. Nevertheless, it could still be a behavioral response of cells to other reduced nutrient availability, such as nitrogen or phosphorus, as these nutrients were eutrophic in silicate-depleted treatments due to silicon limitation. In the control group, however, they were competitively absorbed as the cells proliferated. Nonetheless, it cannot be ruled out that the occurrence of this additional frequency regulation behavior may be specific to certain growth stages of the cells. Further research is needed to determine the specific reasons behind this phenomenon.

Interestingly, the dominant frequency of sinking regulation in both the the control and silicate-depletion conditions ranged from 0.13-0.50 Hz (equivalent to a period of 2 s to 8 s, Fig. 6). The similarity in the dominant frequency of sinking regulation between the control and silicate-depletion conditions suggests that the physiological mechanisms of sinking regulation were the same for both. We currently lack any reports on the frequency of specific physiological mechanisms in diatoms. However, previous studies have reported changes in the electrochemical characteristics of the large diatom *C. wailesii* within a frequency range of 0.01-10 Hz and these changes are believed to be associated with the establishment of electrochemical gradients through ion flux and osmolyte regulation in diatoms (Boyd & Gradmann 1999, Gradmann & Boyd 1999, Boyd & Gradmann 2002). Additionally, researchers conducted on other cell types have identified a frequency range of 0.13-0.50 Hz probably encompasses various physiological mechanisms such as the opening and closing of ion channels, activation and inhibition of intracellular signalling pathways, and regulation of cell morphology and structure through motor proteins and the cytoskeleton (Neher & Sakmann 1976, Findlay & Coleman 1983, Soderling & Beavo 2000, Pollard & Borisy 2003, Demidchik et al. 2018). These mechanisms not only enable cells to adapt to their surrounding environments and respond to external stimuli but also suggest a potential role in the regulation of sinking behavior in large diatoms (Gemmell et al. 2016, Du Clos et al. 2019, Lavoie & Raven 2020, Du Clos et al. 2021). From a time-frequency domain perspective, the frequency or period of regulation in the sinking behavior of *P. hardmaniana* may be influenced by these mechanisms.

## 5. Conclusion

*P. hardmaniana* regulates its sinking behavior by altering the proportions of different instantaneous speed occurrences. The observed regulation of sinking speed was intermittent. The frequency of regulation was 0.13-0.50 Hz (equivalent to a period of 2-8 s) in both control and silicate-depleted conditions. In the time domain, the response of *P. hardmaniana* to silicate depletion, as reflected in both the mean sinking speed and instantaneous sinking speed, exhibited variations over the observation period. However, in the time-frequency domain, *P. hardmaniana* demonstrated an increased regulatory event in response to silicate depletion during the study period. In addition, the oscillation power of sinking regulation was found to be influenced by the physiological state of *P. hardmaniana*, with a poorer physiological state resulting in lower oscillation power of regulation.

